# Budgerigars adopt robust, but idiosyncratic flight paths

**DOI:** 10.1101/598680

**Authors:** Debajyoti Karmaker, Ingo Schiffner, Mandyam V. Srinivasan

## Abstract

We have investigated the paths taken by Budgerigars while flying in a tunnel. The preferred flight trajectories of nine Budgerigars (*Melopsittacus undulatus*) were reconstructed in 3D from high speed stereo videography of their flights in an obstacle-free tunnel. Individual birds displayed highly idiosyncratic flight trajectories that were consistent from flight to flight over the course of several months. We then investigated the robustness of each bird’s trajectory by interposing a disk-shaped obstacle in its preferred flight path. We found that each bird continued to fly along its preferred trajectory up to a point very close to the obstacle before veering away rapidly, making a minimal deviation to avoid a collision, and subsequently returning to its original path. Thus, *Budgerigars* show a high propensity to stick to their individual, preferred flight paths even when confronted with a clearly visible obstacle, and do not adopt a substantially different, safer route. Detailed analysis of the last-minute avoidance manoeuvre suggests that a collision is avoided by restricting the magnitude of the optic flow generated by the obstacle to a maximum value of about 700 deg/sec. The robust preference for idiosyncratic flight paths, and the tendency to pass obstacles by flying above them, provide new insights into the strategies that underpin obstacle avoidance in birds. It could also have wide-ranging implications for conservation efforts to mitigate collisions of birds with man-made obstacles – especially obstacles that are poorly visible, such as wind turbines or buildings with glass facades. Our findings indicate that care needs to be exercised to ensure that newly planned structures are not located near major bird flyways, wherever possible, and to ensure that the positioning takes into consideration the cues and behaviours that birds use to avoid such obstacles.

## 1. Introduction

Recently, there has been growing interest in understanding how birds cope with the challenges of short-range navigation and guidance. Some aspects of visually-guided flight are now beginning to be investigated – such as regulation of flight speed [Schiffner and Srinivasan, 2016], flight between obstacles [Williams and Biewener, 2015, Bhagavatula et al., 2011], collision avoidance [Lin et al., 2014, Schiffner et al., 2016] choice of landing locations [Bhagavatula et al., 2009], flight through narrow apertures [Vo et al., 2016], and body awareness [Schiffner et al., 2014]. However, this challenging area of research is still in its infancy. Typically, these experiments have been conducted in relatively small indoor environments, to facilitate precise experimental control and enable recording and 3D reconstruction of the birds’ flights using multiple high-speed cameras.

Here we investigate the paths taken by birds when they fly in a tunnel whose cross section is large enough to permit a variety of trajectories while moving from one end of the tunnel to the other. Firstly, do the birds exhibit a preferred flight path while flying in a tunnel? If so, does this preference persist with the passage of time? Secondly, do all birds use the same flight trajectory, or does the preferred trajectory vary from bird to bird? Thirdly, if an obstacle is placed in a bird’s preferred path, does it switch to an entirely different flight path, or try to retain its originally preferred flight path by making just a brief detour around the obstacle? The answers should not only provide insights into obstacle avoidance in bird flight, but may also have implications for the siting of structures such as wind turbines and buildings.

## 2. Materials and Methods

### 2.1. Ethics Statement

All experiments were carried out in accordance with protocols approved by the Australian Law on the protection and welfare of laboratory animals, and also by the Animal Experimentation Ethics Committees of the University of Queensland, Brisbane, Australia.

### 2.2. Subjects

English adult Budgerigars, *Melopsittacus undulatus* – four birds, approximately 6–8 years old [*Drongo, Four, Nemo*, and *Two*] – together with five wild-type adult Budgerigars, approximately 2–3 years old [*Halley, Antares, Algol, Pluto*, and *Keppler*] served as subjects for the experiments. The birds were purchased from various local breeders at the age of approximately one month and were housed in a communal mesh walled semi-outdoor aviary at The University of Queensland’s Pinjarra Hills field station. The aviary measured 4 m in length, 2 m in width and 2.2 m in height, and provided a natural diurnal light cycle. The birds also had access, through a window, to a climate-controlled indoor room in an adjoining building, allowing protection from inclement weather. When participating in flight trials (about two to three times a week), the birds were moved to an experimental flight tunnel located at the same station. During the experiments birds were kept in the tunnel in groups of up to four in small cages (47 × 34.5 × 82 cm) for a duration of not more than seven hours per day. After the completion of each day’s experiment, the birds were returned to the aviary.

### 2.3. Experimental Configuration

The experiments took place indoors in a tunnel 25 m long, 1.4 m wide and 2.50 m high, with white side walls, a grey floor and a meshed ceiling. Both ends of the tunnel were covered with a white curtain, to enhance the visual contrast of the bird and facilitate its detection and tracking in the video images (see details below).

Each of the birds was initially trained to take off from a perch at one end of the tunnel (a, Figure 1), fly through the tunnel (with or without an obstacle, depending upon the experiment), and land on a bird cage (1.2 m high) at the other end of the tunnel (b, Figure 1), for five times before video recording of their flights was commenced. The take-off perch was hand-held by an experimenter, and a slow rotation of the perch induced the bird to take off.

**Figure 1:**
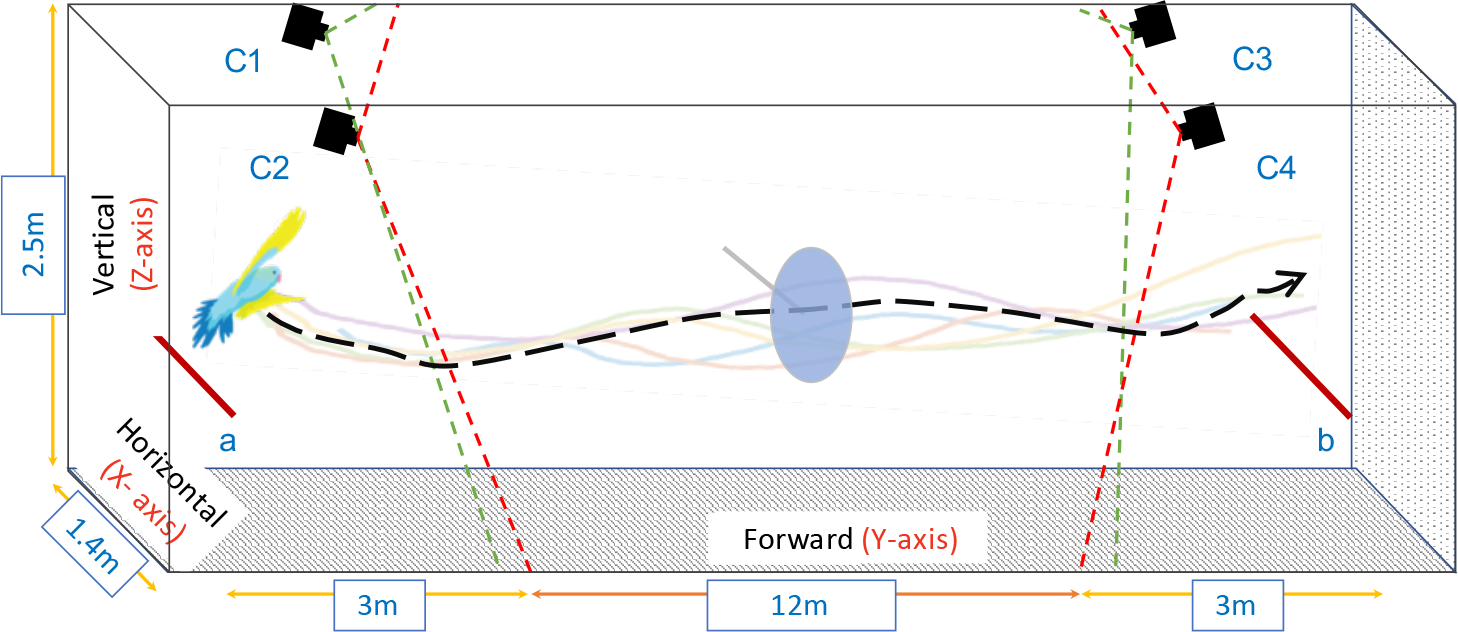
Experimental setup. Birds were trained to fly from point a to b, and from b to a. The mean preferred path for each bird (dashed black line) was calculated by analysing 5 flights (color lines). An obstacle (a blue disk, diameter 41cm) was then placed in the tunnel to obstruct the preferred path of each bird, and the flights were re-filmed. The flights were recorded using four synchronised high-speed cameras (C1-C4), mounted on the side walls. The figure is not to scale.

Two different scenarios were presented to the birds: (i) obstacle-free flights (rounds 1 and 2); and (ii) obstacle flights. The first obstacle-free scenario (round 1) served as a baseline for establishing each bird’s preferred flight path. The second obstacle-free scenario (round 2) was conducted eight months later, to check whether the birds retained their originally preferred path.

The obstacle flights were commenced 4 days after the completion of the second obstacle-free scenario. In these flights each bird was confronted with an obstacle (a disk 41 cm in diameter; made from thin blue cardboard, supported by a thin horizontal dowel), placed in the bird’s preferred flight path (as defined by the mean position of the flight path over a segment ±1 m from the disk along the longitudinal axis of the tunnel). As will become evident in the Results section, individual birds displayed consistent, but idiosyncratic flight paths. For each bird and flight direction, the disk was positioned according to the bird’s mean flight position during the obstacle-free flights in that direction (a to b, or b to a). However, from the perspective of each bird the position of the disk was very similar for the two flight directions.

Nine birds (*Drongo, Four, Nemo, Two, Halley, Antares, Algol, Pluto*, and *Keppler*) were tested individually in each of the three scenarios. For the first round of obstacle-free flights, after the training phase we recorded 5 flights from each side of the tunnel for each bird, summing up to 90 flights in total. For the second round of obstacle-free flights, we recorded 2 flights from each side for each bird, summing up to 36 flights. Finally, for the obstacle flights, we recorded 5 flights from each side for each bird, summing up to 90 flights. Altogether, 216 flights were recorded in the course of the study.

### 2.4. Recording

The flights were recorded using a network of four synchronised Emergent HS-4000 ultra-high speed 10GigE 4-mega-pixel cameras (C1, C2, C3 and C4 in Figure 1). The cameras were mounted on the two side walls at both ends of the tunnel at a height 2.2 m, as illustrated in Figure 1. Each camera was equipped with a narrow-angle lens (12 mm focal length) to maximise the image resolution within the region of interest. Each flight yielded four synchronised image sequences. Flights were recorded at 100 frames per second, which provided adequate temporal resolution for detecting and tracking the birds.

### 2.5. Reconstruction of 3D Trajectories

For the reconstruction of the birds’ flight trajectories in 3D, cameras C1 and C2 were used for flights from point b to a, and cameras C3 and C4 for flights from a to b (see Figure 1). This selection of camera pairs enabled reliable detection of the bird from the take off point and ensured consistent tracking of the location of the bird’s head in each video frame, regardless of the flight direction. The videos of each flight from the two cameras were initially processed using a bird detection and tracking algorithm [Karmaker et al., 2018] to locate the centroid of the bird’s head location in each frame, and the positions in stereo frames were then combined to generate the 3D trajectory of the bird’s flight.

Stereo calibration of the cameras was carried out using a reference checker-board pattern (check size 82.5 mm) in association with the Stereo Camera Calibrator Toolbox for Matlab (MathWorks^®^). This procedure delivered the calibration parameters for each camera, and also determined the precise 3D position and orientation of one camera relative to the other. The mean overall reprojection error in our experimental configuration was estimated to be 0.19 pixels, by using the error estimator built into the Stereo Camera Calibrator Toolbox.

We computed the 3D flight profiles of the birds, and plotted the variation of the radial distance of the bird with respect to the obstacle, across each flight. We also computed the profile of the optic flow generated by the obstacle in the visual system of the experimental bird during its flight through the tunnel. To facilitate visualisation and interpretation, these profiles were plotted on a distance scale (along the Y axis) by appropriate interpolation of the respective time functions.

### 2.6. Statistical Evaluation

In order to test whether individual birds followed preferred flight paths, we performed a nearest-neighbour evaluation for each set of flights and compared each set to a set comprising an equal number of flights from randomly selected individuals, using the Wilcoxon signed rank test for paired samples. For further statistical analysis of the general flight profiles, for example comparing the horizontal and vertical positions of the birds at the point of crossing the obstacle across the three scenarios, we used the Aligned Rank Transformed (ART) ANOVA test using a linear mixed effects model [Wobbrock et al., 2011], with the Experimental Condition and Direction of Release as fixed effects and the Bird as a random effect. This type of ANOVA does not require the data to be normally distributed; we did, however, confirm that the variances of the samples were homogeneous. For post hoc comparison, we employed least squared means using the Tukey method for multiple comparisons.

In order to test whether the birds displayed a preference for passing the obstacle (disk) in a particular direction (for example, above or below it, or to its right or left), we used the methods of circular statistics. The direction in which a bird (bird *i*) cleared the disk was quantified by computing the mean direction over its 10 flights, and representing this mean direction by a unit vector 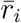. The mean clearing direction, 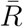, averaged over the nine birds, was then computed as the average of the nine unit vectors (one per bird) [Batschelet, 1981]: 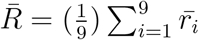

The direction of 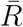 indicates the overall mean clearing direction (averaged over all 9 birds), and the length of 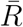, (| *R* |), provides a measure of the coherence in the clearing direction across the nine birds: |*R*| has a maximum value of 1 when all birds display exactly the same clearing direction, and a minimum value of 0 when the clearing directions are distributed randomly [Batschelet, 1981]. The Rayleigh test [Batschelet, 1981] was used to obtain a *P* value for ascertaining whether the distribution of clearing directions, across all birds, was significantly different from random.

## 3. Results

### 3.1. Obstacle-free Flights

In the obstacle-free flights (round 1) we noticed a high propensity for individual birds to stick to specific flight paths. This is illustrated in Figure 2, which shows the mean horizontal position (Figure 2a) and the mean vertical position (Figure 2b) for each bird, averaged over its entire flight. To verify the tendency of each bird to fly along a specific path, we used a nearest neighbour approach to compare the average nearest neighbour distance for each set of flights of a given bird with a set of flights from randomly selected birds. We found that each set of flights for a given bird had a significantly smaller nearest neighbour distance compared to a set of flights selected randomly across the population (see row 1, Table 1), indicating that each bird indeed tended to stick to its preferred flight path.

**Figure 2:**
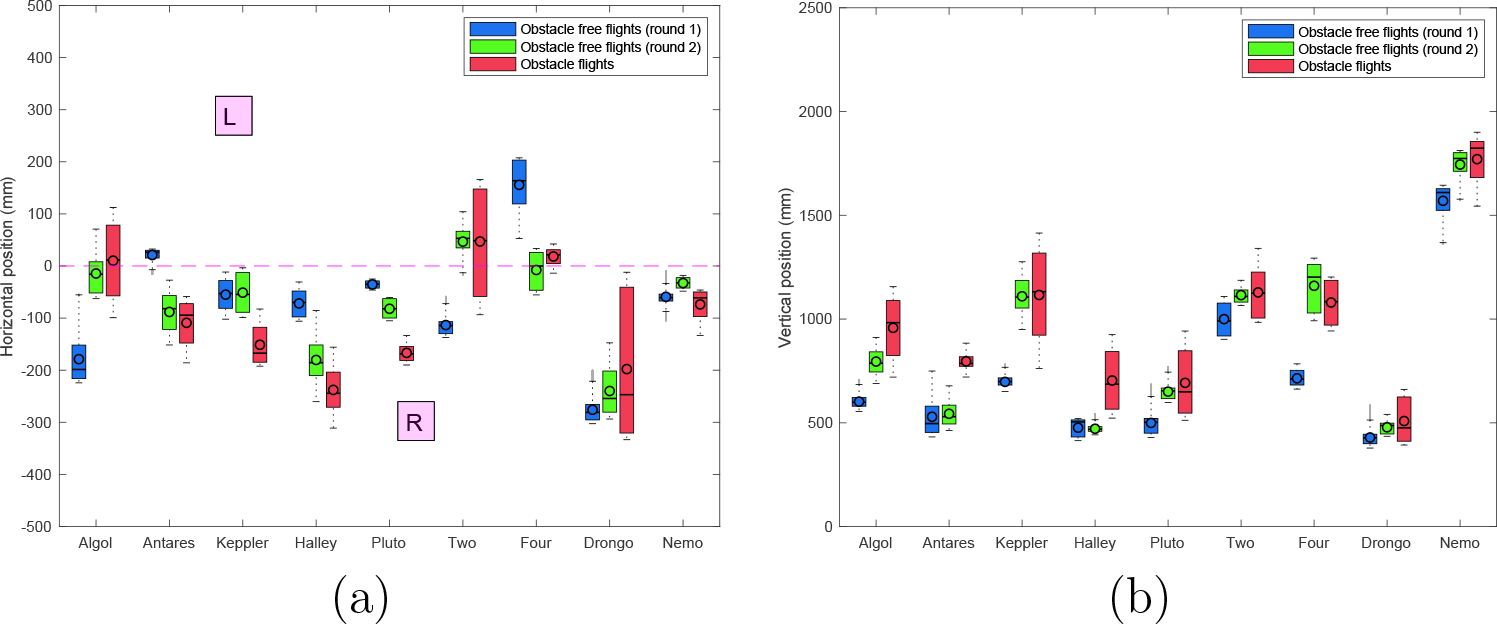
Box-and-whisker plot showing the average relative horizontal position (a) and altitude (b) of each bird - averaged over the entire flight - in each of the three experimental scenarios. The black circles represent the mean values and the black dashes represent the median values. The whiskers indicate the variability outside the upper and lower quartiles. Data not included between the whiskers, i.e. outliers, are represented as grey lines. In (a) the pink dashed line depicts the midline of the tunnel. R and L denote positions to the right and left of the tunnel, respectively, from the bird’s point of view.

**Table 1:** Grand median of the average nearest-neighbour distance (mm) for each set of flights and the respective randomised set of flights.

|  | Actual flights<br>grand median | Randomised flights<br>grand median | Wilcoxon signed<br>rank test |
| --- | --- | --- | --- |
| Obstacle-free flights (round 1) | 233 | 436 | $V = 0; p = 7.629e^{-06}$ |
| Obstacle-free flights (round 2) | 212 | 471 | $V = 22; p = 0.004$ |
| Obstacle flights | 236 | 513 | $V = 0; p = 7.629e^{-06}$ |

The tendency of each bird to fly a fixed path was evident for flights in both directions in the obstacle-free tunnel. This is illustrated in Figure 3, which shows the mean position of each bird in the cross section of the tunnel (from the bird’s viewpoint), during its flights in the forward and the reverse directions in round 1. The mean positions are very similar for the flights in the two directions – the average distance between these positions is 19.89 ± 9.54 cm (SD). This further confirms the robustness of each bird’s preferred path.

**Figure 3:**
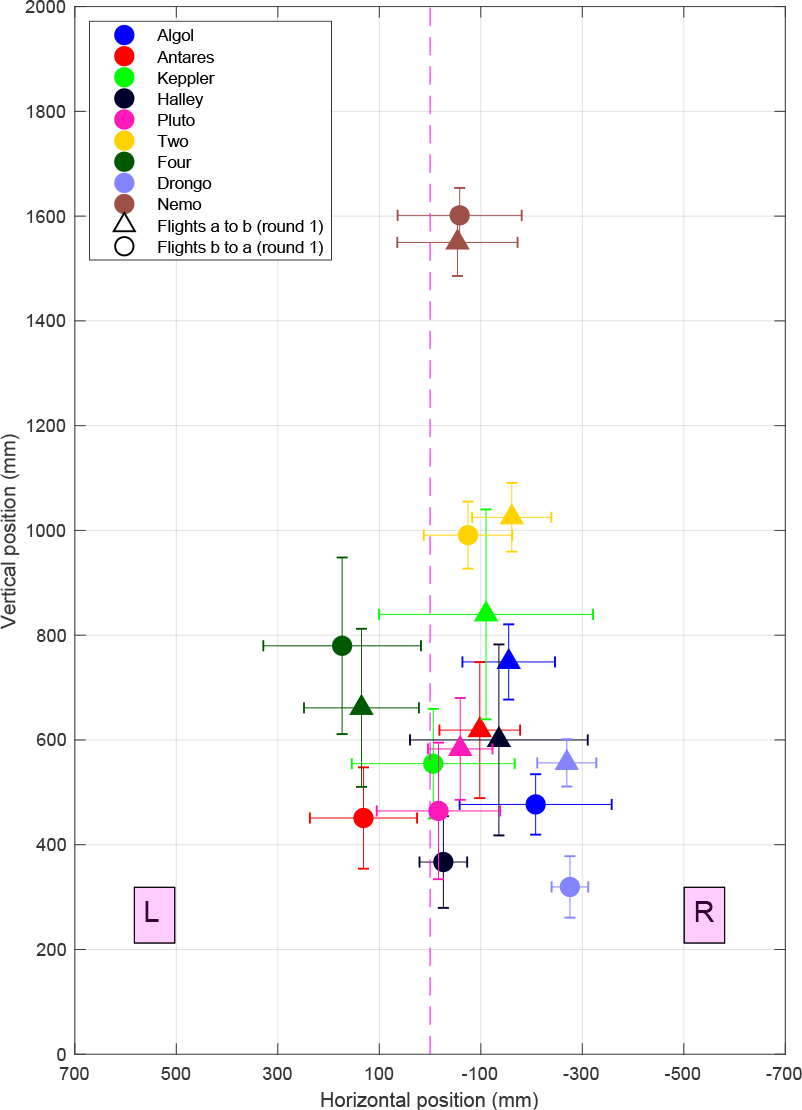
Mean position of each bird during the flights forward vs reverse directions in round 1.

In round 2 of the obstacle-free flights there was again a strong tendency for individual birds to stick to specific flight paths (see Figures 2a, 2b). Again, each set of flights for a given bird had a significantly smaller nearest-neighbour distance compared to a set of flights selected randomly across the population (see row 2, Table 1), indicating that each bird maintained a robust flight path.

A comparison of the results obtained for the two obstacle-free scenarios reveals that, in all of the experimental scenarios there is a slight tendency for the birds to fly slightly to their right of the mid-line of the tunnel (irrespective of the flight direction). However, for the most part (and with a few exceptions), there is no major change in the overall mean horizontal position or the overall mean vertical position of each bird’s preferred trajectory between rounds 1 and 2. This is confirmed in the plot of Figure 4a, which compares the mean position of each bird’s trajectory in the cross section of the tunnel, across the obstacle-free scenarios (rounds 1 and 2). While three birds (*Algol*, *Four* and *Keppler*) changed their mean positions between the two rounds of obstacle-free flights – mostly in the vertical plane – the other six birds maintained approximately the same average positions (see Figure 4a, squares vs circles). Thus, the majority of the birds maintained their preferred trajectories over the 8-month interval between the experiments conducted in rounds 1 and 2. A detailed statistical comparison of these results is given later below.

**Figure 4:**
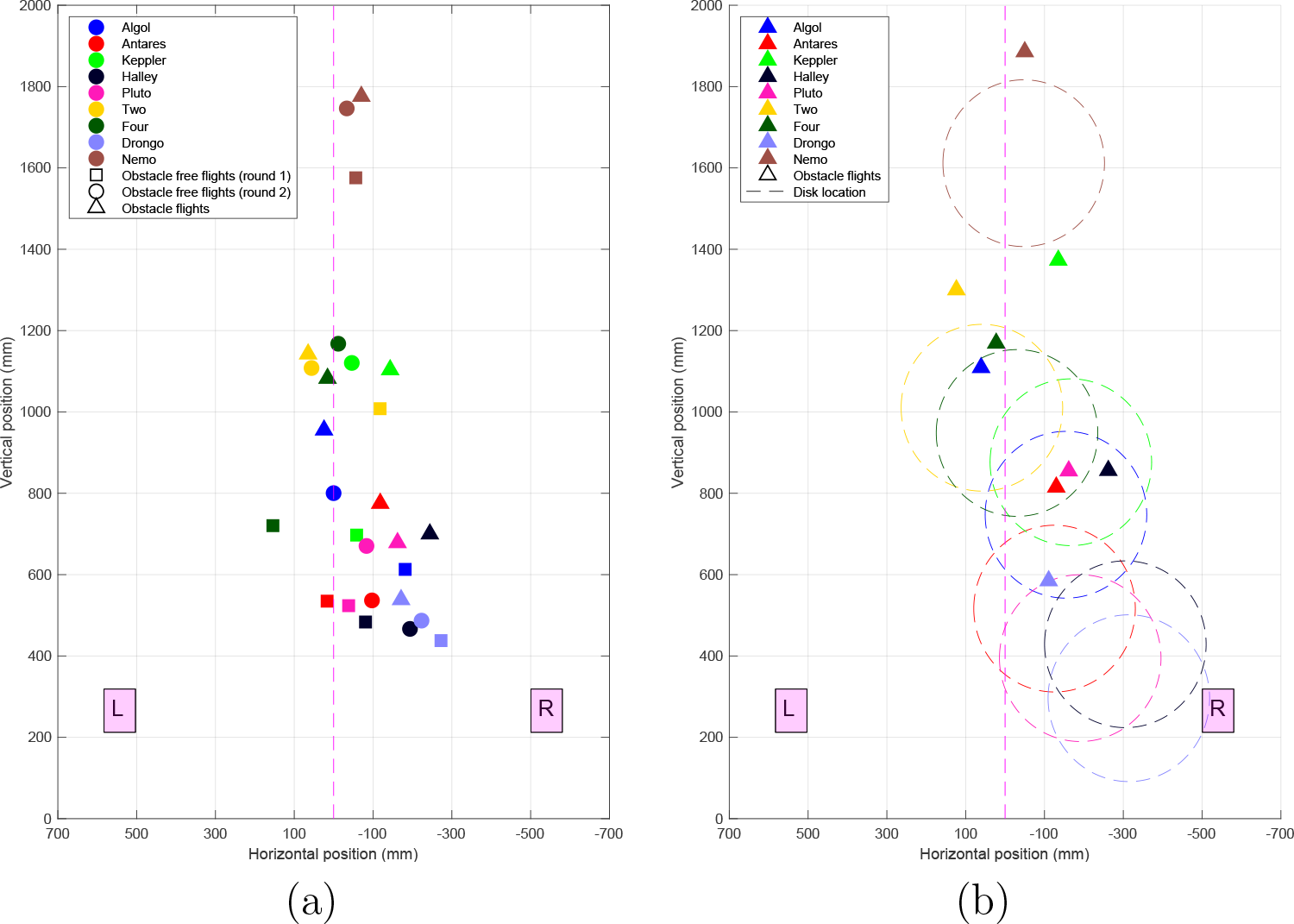
(a) Mean position of each bird - averaged over the entire flight – for three experimental scenarios. Each color represents an individual bird. The squares and circles denote the obstacle-free flights (rounds 1 and 2, respectively), and the triangles denote the obstacle flights. (b) Comparison of the mean position of each bird in the cross section of the tunnel in the obstacle-free scenario (round 2) and in the obstacle scenario. In each case the mean position represents an average over a 2 m flight segment spanning the obstacle (±1 m from the obstacle). The dashed circles represent the location of the obstacle for each bird. The centre of the circle represents the (local) mean position of each bird in round 2 of the obstacle-free flights. The dashed vertical line (pink) depicts the centre of the tunnel. R and L denote positions to the right and left of the tunnel, respectively, from the bird’s point of view.

### 3.2. Obstacle Flights

The question then arises: How does a bird behave if an obstacle is interposed in its preferred flight path? To examine this, we recorded 10 flights of each bird when an obstacle, consisting of a 41 cm diameter disk, was positioned with its centre located at the mean position of the bird’s trajectories in round 2 of the obstacle-free scenario, as measured over a 2 m flight segment spanning a range ±1 m from the position at which the disk was located in the obstacle tests. Figure 5 shows the position of each bird in the cross section of the tunnel in round 2 of the obstacle-free scenario, relative to the position at which the disk was placed in the subsequent obstacle tests. It is clear that the mean position of each bird lies within the projected cross section of the disk, at distances of up to 4 m ahead of the position at which the disk was placed in the subsequent obstacle tests. Thus, the disk was squarely in the flight path of each bird, as measured in the obstacle-free scenario.

**Figure 5:**
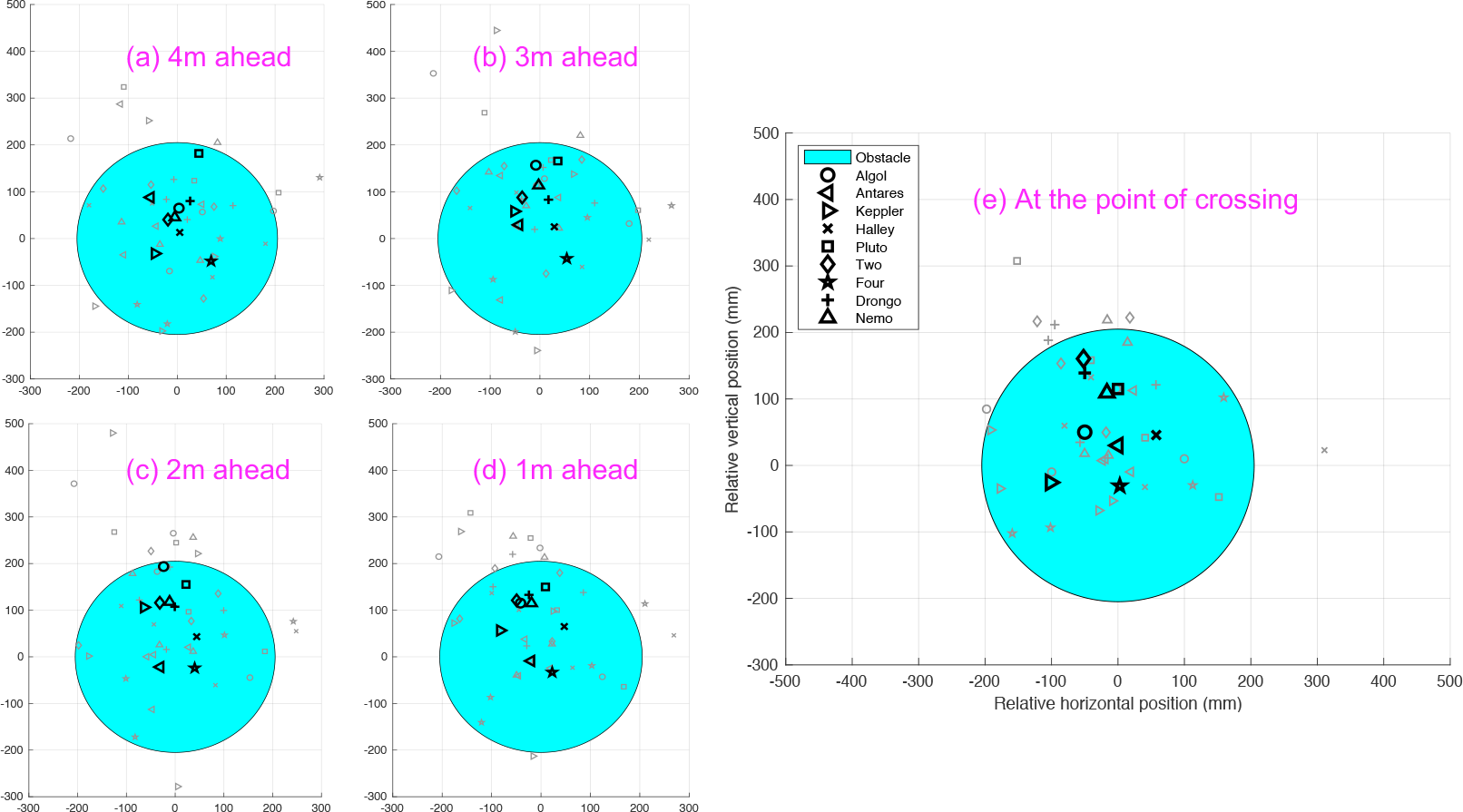
Position of each bird in the cross section of the tunnel for each of its 4 flights in round 2 of the obstacle-free scenario (small symbols), relative to the position at which the disk was placed in the subsequent obstacle tests, at different axial distances from the disk, as well as at the point of crossing. The large symbols represent the mean position of each bird, averaged over 4 flights.

The obstacle tests were commenced 4 days after the tests in round 2 of the obstacle-free scenario had been completed. Hence, it was very unlikely that the birds would have changed their flight path preferences substantially during this short interval.

Figure 4b compares the mean positions of each bird in the obstacle scenario and in the obstacle-free scenario (round 2), over a 2 m flight segment spanning the position of the obstacle (±1 m from the obstacle). The dashed circles indicate the size and position of the obstacle (disk) for each bird. It is evident that each bird avoids the disk by flying above it.

The avoidance manoeuvre is analysed in greater detail in Figure 6, which shows scatter-plots of the radial positions of the birds relative to the obstacle when they were at distances of 4 m, 3 m, 2 m, and 1 m ahead of the obstacle, and at the point of crossing the obstacle. Even in the presence of the obstacle, the birds maintained their preferred paths up to a point 3 m ahead of the obstacle (Figure 6a), beyond which they started to veer away progressively (Figures 6b,c,d), eventually clearing the obstacle by flying above it (Figure 6e). The veering response to avoid the obstacle is quantified by the magnitude and direction of the mean vector 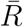 (Figure 6e), computed as described in ‘Materials and Methods’. 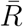 is directed upwards, has a magnitude close to 1.0, and is highly significant (*P* < 0.00002), indicating that the obstacle is avoided by flying above it.

**Figure 6:**
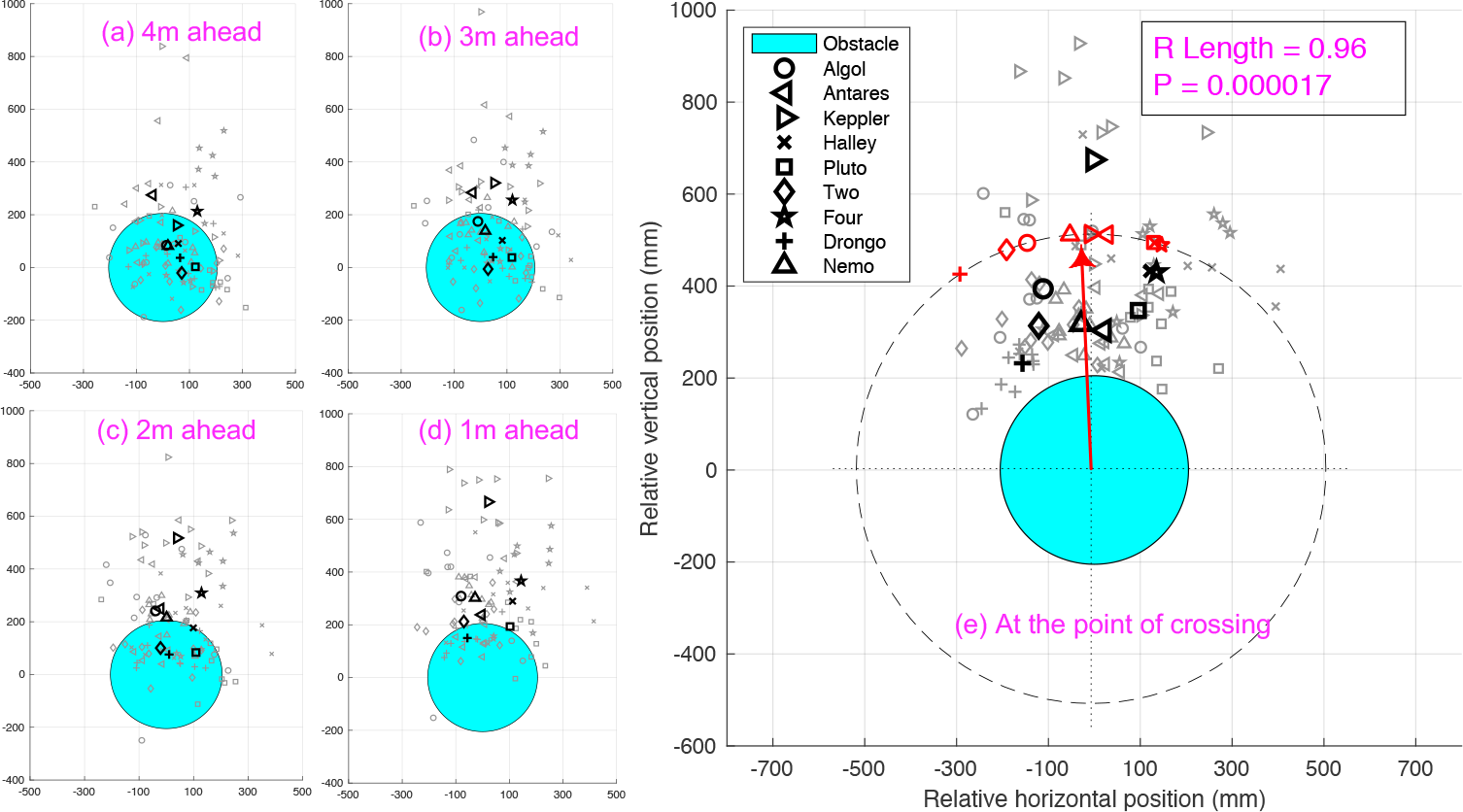
Horizontal and vertical positions of the birds relative to the obstacle, at distances of 4 m (a), 3 m (b), 2 m (c), and 1 m (d), ahead of the obstacle, and at the point of crossing the obstacle (e). The blue circle depicts the location of the obstacle relative to the birds’ flights. The small black symbols represent the positions of the flights of individual birds (10 per bird, totalling 90 flights), and the large black symbols represent the mean position for each bird. In (e) the red symbols represent the direction of the mean clearing vector 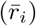 for each bird, and the arrow shows the overall clearing vector 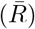, averaged over all birds. The dashed circle represents the largest possible magnitude of the clearing vector (1.0). Details in ‘Materials and Methods’.

Figure 7 examines the obstacle avoidance response in another way, by charting the percentage of flights that are on a collision course with the obstacle, as a function of the distance from the obstacle. This is done by determining, at each distance, the number of flights in which the bird is positioned within the volume of the axial cylinder projected by the disk (red columns), and outside it (grey columns), as can be visualised in Figure 6. Figure 7 indicates a substantial decrease in the percentage of collision-directed flights between 3 m and 2 m, suggesting that collision avoidance commences at a distance of ~ 2.5 m from the obstacle. The blue/green columns in Figure 7 show the corresponding numbers for round 2 of the obstacle-free scenario, for a ‘virtual’ disk placed at the position where it was located in the obstacle tests. In this case the percentage of flights within the virtual disk does not change substantially – it is more or less constant at a high level (70% − 75%), demonstrating that, in the obstacle tests, the disk was indeed placed directly in each bird’s preferred flight path, as measured in round 2 of the obstacle-free scenario.

**Figure 7:**
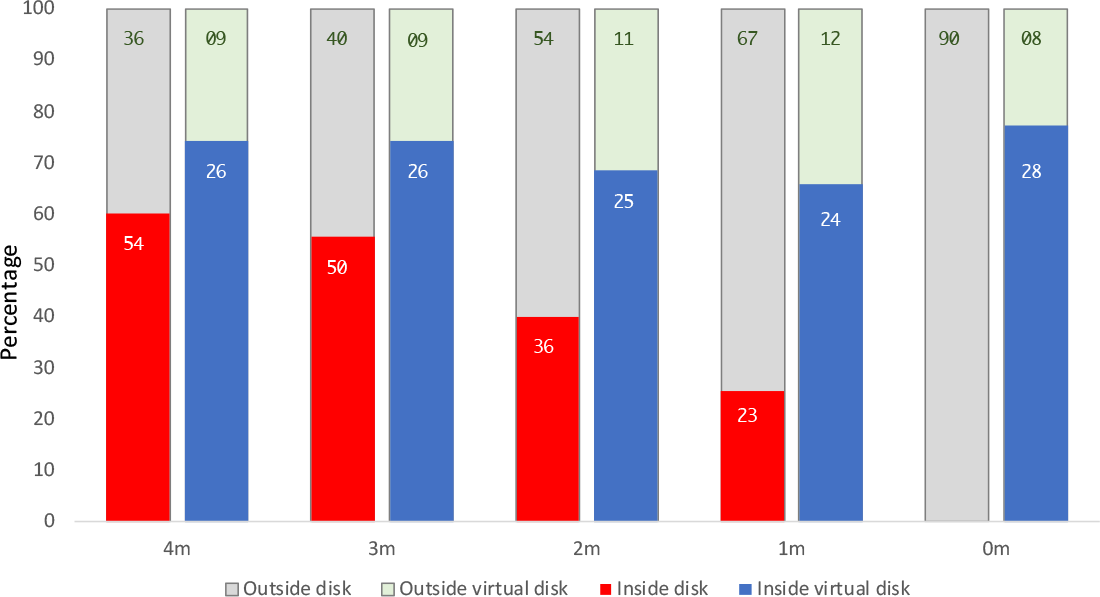
Percentage of flights that are on a collision course with the disk, at different axial distances from the disk. The red/grey columns show the numbers of flights that are within the projected area of the disk (i.e. on a collision course) or outside the projected area (i.e. not on a collision course). The blue/green columns show the corresponding figures for round 2 of the obstacle-free scenario, for a ‘virtual’ disk placed at the position where it was located in the obstacle tests. Each bird performed 10 flights in the obstacle tests (totalling 90 flights), and 4 flights in round 2 of the obstacle-free scenario (totalling 36 flights). Note that in round 2 at 4 m and 3 m the total number of flights is 35 as *Pluto*, during one flight was recorded only after the 3 m mark.

Finally, a nearest-neighbour analysis of the obstacle flights reveals that, as in the obstacle-free scenarios (rounds 1 and 2), each set of flights for a given bird has a significantly smaller nearest-neighbour distance compared to a set of flights selected randomly across the population (see row 3, Table 1), indicating that each bird tends to stick largely to its preferred flight path when avoiding the obstacle.

### 3.3. Control of Radial Separation from the Obstacle

Next, we investigated in greater detail how the birds controlled their radial separation from the obstacle (disk) during their flight. Figure 8 shows the profiles of the mean radial separation (defined as the radial distance of the bird’s head from the centre of the disk) for the obstacle flights (orange circles) and for round 2 of the obstacle-free scenario (blue circles). The two profiles are similar up to a point 3 m away from the obstacle. At distances closer to the disk, the birds begin to veer away from the disk when it is present, achieving a maximum radial separation of about 500 mm slightly beyond the point of crossing the disk. This radial separation represents a distance of 295 mm from the edge of the disk, which has a radius of 205 mm.

Figure 9 shows the difference between the mean radial distance profiles for the obstacle-free flights (round 2) and the obstacle flights shown in Figure 8, normalised to a value of 100. During the initial phase (−6 m to −2 m, highlighted by the yellow rectangle) the difference between the two profiles is less than 10%, indicating that, during the initial part of their flight, the birds maintain their preferred trajectories regardless of the presence or absence of an obstacle in their path. In the presence of an obstacle, the birds begin to deviate from their preferred trajectory only when they are about 2.5 m away from the obstacle. After passing the obstacle the birds tend to return to the originally preferred obstacle-free trajectories, as measured in round 2.

**Figure 8:**
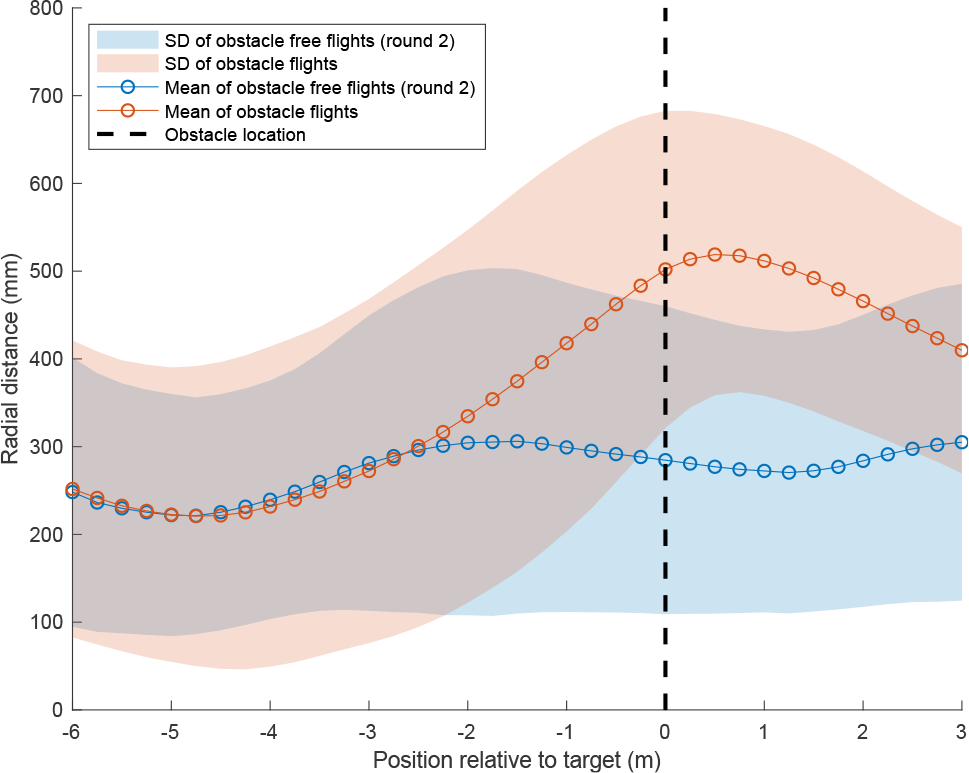
Mean radial distance profiles and standard deviation (SD) for the obstacle-free flights (round 2) and the obstacle flights. The vertical dashed line depicts the position of the obstacle.

**Figure 9:**
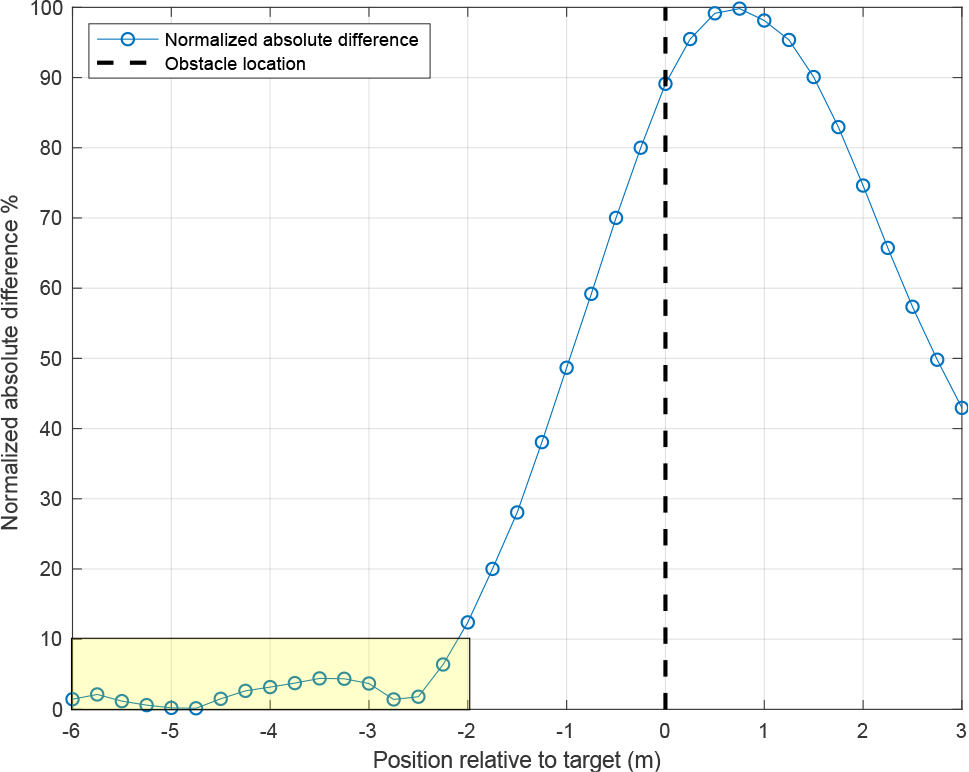
Normalised absolute difference between the mean radial distance profiles for the obstacle-free flights (round 2) and the obstacle flights. The vertical dashed line depicts the position of the obstacle.

We also computed, for the obstacle scenario, the time course of the optic flow (image angular velocity) generated by the the disk as the bird flew past it (Figure 10). The magnitude of the optic flow increases rather sharply as the bird approaches the disk, reaching a maximum close to the point of crossing the disk. It is of interest to note that all of the birds fly past the disk in such a way as to restrict the maximum optic flow to about 700 deg/s. This raises the possibility that, while flying past the disk, the birds are maintaining a safe distance from the the disk by ensuring that the magnitude of the optic flow generated by the disk does not exceed 700 deg/s.

**Figure 10:**
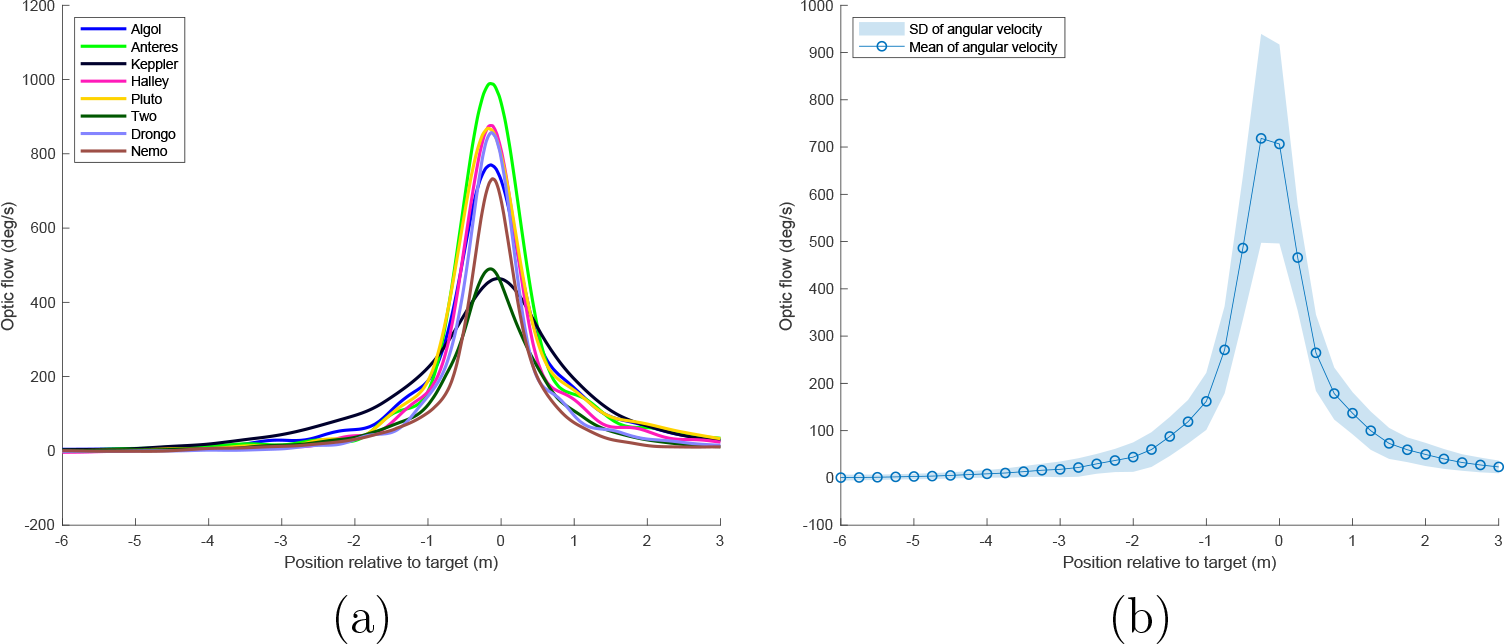
Profiles of optic flow (image angular velocity, in deg/s) generated by the obstacle at each point of the flight. (a): Average profiles for individual birds; (b): Mean profile averaged over all birds.

If the birds are sensing the optic flow generated by the the disk and using it to control the radial distance to the disk, they would need to hold their flight speed constant – only then can the optical flow be directly calibrated in terms of distance. Table 2 compares the coefficients of variation (CV) of the flight speed, radial separation and optic flow. The CVs of the radial distance and the optic flow are significantly different from each other (Wilcoxon signed rank test:V=4; p-value=0.02734). The CV of the flight speed, however, is also significantly lower than that of the radial separation (V=0; p-value=0.003906) and the optic flow (V=1; p-value=0.007812). This indicates that the speed of flight is tightly controlled, as predicted by the optic flow hypothesis.

**Table 2:** Mean flight speed, radial separation and optic flow, and their coefficients of variation at the point of crossing the obstacle for all birds.

|  | Flight speed (m/s) |  |  | Radial separation (mm) |  |  | Optic flow (deg/s) |  |  |
| --- | --- | --- | --- | --- | --- | --- | --- | --- | --- |
|  | MEAN | SD | CV | MEAN | SD | CV | MEAN | SD | CV |
| **Algol** | 7.318 | 0.326 | 0.045 | 566.367 | 163.811 | 0.289 | 729.223 | 172.968 | 0.237 |
| **Antares** | 8.126 | 0.656 | 0.081 | 444.274 | 88.142 | 0.198 | 937.668 | 99.697 | 0.106 |
| **Keppler** | 6.681 | 0.617 | 0.092 | 834.79 | 150.080 | 0.18 | 462.864 | 99.916 | 0.216 |
| **Halley** | 6.537 | 0.479 | 0.073 | 445.554 | 139.934 | 0.314 | 810.672 | 256.826 | 0.317 |
| **Pluto** | 7.338 | 0.514 | 0.070 | 493.724 | 120.348 | 0.244 | 796.696 | 147.532 | 0.185 |
| **Two** | 4.377 | 0.249 | 0.057 | 519.38 | 53.705 | 0.103 | 451.525 | 39.234 | 0.087 |
| **Four** | 4.064 | 0.383 | 0.094 | 595.678 | 238.485 | 0.400 | 404.698 | 118.423 | 0.293 |
| **Drongo** | 5.478 | 0.461 | 0.084 | 368.212 | 75.711 | 0.206 | 788.524 | 122.539 | 0.155 |
| **Nemo** | 4.534 | 0.627 | 0.138 | 343.442 | 62.200 | 0.181 | 675.021 | 81.933 | 0.121 |
| **MEAN** | **6.050** | **0.479** | **0.082** | **512.380** | **121.38** | **0.235** | **672.988** | **126.563** | **0.191** |

Figure 11 shows the mean %CV of the flight speed along the axis of the tunnel at various axial distances from the obstacle. These values are uniformly low, the largest value being 12.2%. Interestingly, the CV decreases as the bird approaches the obstacle and attains it lowest value in the vicinity of the obstacle, indicating tightest control of flight speed in that region. This further supports the optic flow hypothesis.

**Figure 11:**
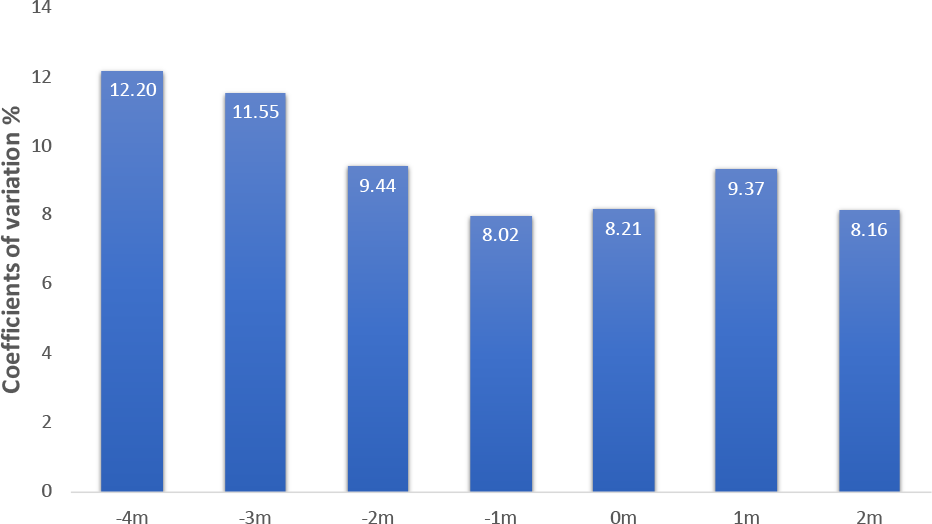
Mean coefficient of variation (%), computed as in Table 2, at various axial distances from the obstacle.

Another prediction of this hypothesis is that, if the birds are regulating the distance to the obstacle by maintaining a constant optic flow, then higher flight speeds should be associated with larger radial distances. Figure 12 shows the correlation between flight speed and the distance to the disk at the point of crossing the disk, for those birds where the correlation was found to be significant. A linear regression on the data reveals that radial distance is indeed correlated positively with speed, as predicted by the optic flow hypothesis. Taken together, the data in Figures 9 and 10 strongly suggest that the birds are keeping a safe distance from the the obstacle by ensuring that the peak optic flow generated by the disk does not exceed a value of about 700 deg/sec.

**Figure 12:**
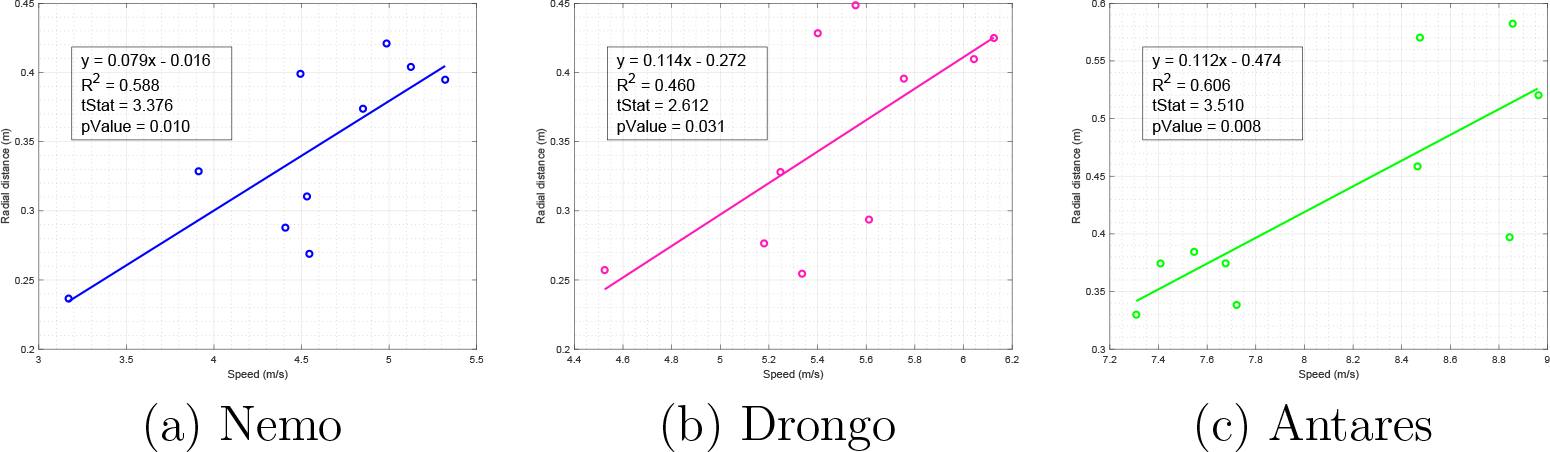
Examples of correlation between flight speed and radial distance to the obstacle at the point of crossing the obstacle for those birds where the correlation is significant.

### 3.4. Recovery After Passing the Obstacle

Figure 9 shows a steep reduction of the radial separation after the birds have passed the obstacle. Although the visual field of the stereo camera system did not extend beyond a distance of 3 m past the obstacle, the radial separation has already dropped to about 40% of its maximum value at this point, and the continued steep downward trend in the radial separation profile strongly suggests that each bird tends to return to its originally preferred trajectory soon after passing the obstacle. This notion is supported by the data in Figure 4a, which compares, for each bird, the mean positions of the flight in the cross section of the tunnel in round 2 of the obstacle-free scenario (circles) and in the obstacle scenario (triangles), averaged over the entire flight. The two mean positions are close together, for each bird. The average separation is 11.76 ± 7.6 cm (SD), and never exceeds 23.88 cm. This small separation suggests that each bird does indeed merge into its originally preferred trajectory after crossing the obstacle, and does not execute a new flight path.

### 3.5. Statistical Meta-Analysis of Flight Trajectories in the Three Scenarios

The flights recorded in the obstacle-free scenario (rounds 1 and 2) and in the obstacle scenario were analysed using ART-ANOVA statistics to look for scenario-dependent effects on the mean horizontal positions and altitudes of the trajectories. The analysis revealed that there was no significant effect of scenario on the mean horizontal position (ART-ANOVA: *F*_2,16_ = 0.55605; *p* = 0.58416). However, there was a significant effect of scenario on the mean vertical position (ART-ANOVA: *F*_2,16_ = 5.2214; *p* = 0.017968). Post hoc comparison revealed that this effect was limited to a change in the mean flight altitude between round 1 of the obstacle-free flights and the obstacle flights (lsmeans: *df* = 16; *t* = 3.210; *p* = 0.0143). There was no significant change in the mean flight altitude between round 2 of the obstacle-free scenario and the obstacle flights (lsmeans: *df* = 16; *t* = 1.284; *p* = 0.4239). The difference between rounds 1 and 2 of the obstacle-free flights did not quite reach statistical significance (lsmeans: *df* = 16; *t* = −1.926; *p* = 0.1636).

In summary, there was no change in the preferred mean horizontal position for each bird, over all three scenarios. There was also no consistent change in the preferred mean altitude. The small differences observed may therefore be the result of minor changes in the height preferences of a few individual birds over the course of the 8 months that had elapsed between the two sets of obstacle-free experiments.

This analysis corroborates our findings that (a) Budgerigars display flight trajectories that are robust, but idiosyncratic; and (b) the introduction of an obstacle in a Budgerigar’s flight path does not alter its trajectory in a major way. The bird largely retains its original trajectory, except for a brief manoeuvre to avoid the obstacle.

## 4. Discussion

We have shown that Budgerigars display a high propensity to stick to individually preferred flight paths, as indicated by the significantly smaller nearest neighbour distance of each set of flights compared to a set of random flights. While our experiments were set in an artificial environment, similar behaviours have been reported for free ranging birds [Biro et al., 2004, Meade et al., 2005]. Like our birds, pigeons, after several releases from the same location, develop a behaviour called route stereotypy, where a given individual retreads the path it has previously taken. This behaviour is generally associated with pigeons navigating to their goal using a simple strategy known as ‘piloting’, which involves moving from one familiar landmark to the next, or using landmarks in more complex ways to guide the flight trajectory. The difference in our experiment is that the observed stereotypical behaviour is likely embedded in an intrinsic motor pattern that is not associated with external landmarks – given that the environment presented to our birds was largely devoid of landmarks, the flights were short, and the goal was always in view.

This raises the question as to why the birds show this kind of stereotypical behaviour. It is important to note that the individual birds’ preference was consistent over small time scales, and also remained largely consistent over larger time scales (8 months). But, with no alterations made to the experimental setup, the most likely reason for the slight changes in preference displayed by some of the birds (over 8 months) would have to do with the bird itself. As to why only some birds change their preference, we can only guess. If we assume – and this is currently our best guess – that the flight path preference is associated with the position that the bird would take up if it were flying in a flock, then the reason for a change in preference could be related to a change in the hierarchical order of the birds. Unfortunately, we could not test this possibility as it was not feasible, given the constraints of the tunnel, to release all birds simultaneously. However, this would be an interesting and worthwhile direction of further research.

Our results (Figures 2, 4 and 6) reveal that Budgerigars tend to avoid a small obstacle by flying above it, rather than below it, or to either side. In this context, it is important (and interesting) to note that some of the birds (such as *Nemo, Two, Four* and *Keppler*) flew at a rather high altitude (see Figure 2b), thus requiring the obstacle to be placed at large heights – close to the ceiling of the tunnel – to obstruct their flight paths. Despite this, and despite the fact that there was a large clear space under the obstacle, even these birds avoided the obstacle by flying above it, rather than below it. Thus, all of the 9 birds avoided the obstacle by flying above it when it was placed in their flight path, regardless of the height of the obstacle. Another important aspect is that the birds maintained a distance of around 30 cm (see Figure 8) from the boundary of the obstacle at the point of crossing it, which is roughly equal to the average wing span (30 cm) of a Budgerigar. This separation should ensure avoidance of the obstacle with a minimal deviation from the preferred flight trajectory – which speaks volumes for the body awareness of Budgerigars. This feature of body awareness has also been demonstrated in earlier studies investigating flights of Budgerigars through narrow vertical apertures [Vo et al., 2016, Schiffner et al., 2014].

The most surprising observation, however, is that Budgerigars show a strong tendency to retain their preferred flight path even when an obstacle is introduced in the path. The obstacle is avoided with a minimal and potentially dangerous last-minute manoeuvre. Our data, analysed in two different ways (see Figure 7 and Figure 9) infer that the avoidance manoeuvre commences at a distance of ~ 2.5 m. The average flight speed of about 6 m/s (as shown in Table 2) implies that the initiation of the avoidance occurs less than half a second before the obstacle is encountered.

Bhagavatula et al. [Bhagavatula et al., 2011] demonstrated how Budgerigars (*Melopsittacus undulatus*) navigate through the middle of a tunnel by balancing the magnitude of optic flow (the speed of image motion) experienced by the bird’s eyes. Our findings suggest that the birds using the same cue (optic flow) to regulate their flight while avoiding a stationary disk like obstacle. They are accomplishing this avoidance behaviour by keeping a safe distance from the obstacle by ensuring that the peak optic flow generated by the center of gravity of the disk does not exceed a value of about 700 deg/sec. It is possible that the birds are sensing the largest magnitude of the optic flow generated by disk – that by the nearest margin – boundary, rather than the flow generated by its center of gravity. However, this does not negate the optic flow hypothesis, as it would simply scale up the optic flow threshold.

It is noteworthy that the individual birds display a remarkably constant speed throughout their flight. This greatly simplifies the use of cues based on optic flow to compute range, as the distance to an obstacle can then be directly calibrated in terms of the magnitude of the optic flow induced in the eye [Altshuler and Srinivasan, 2018].

It is very unlikely that the reason for the late avoidance response is due to a lack of visibility of the obstacle. When the bird begins to veer away from the disk at a distance of 2.5m, the disk would subtend a visual angle of 9.4 deg, and present a high contrast against the white background at the end of the tunnel. Earlier experiments, investigating the ability of Budgerigars to find and land on high-contrast disks of a similar size, confirm that such objects are clearly visible to the birds [Bhagavatula et al., 2009].

These findings have potentially wide-ranging implications for decisions about the location and design of wind-farms, and other obstacles with which birds are prone to collide. When choosing potential locations for new wind-farms, care has to be taken to position them away from major bird flyways, wherever possible, and to ensure that the positioning takes into consideration the strategies that birds are likely to use to avoid such obstacles.

## 5. Supporting Information

Videos showing that Budgerigars show a strong tendency to retain their preferred flight path even after several trails with the obstacle placed in their respective flight path. Available at: https://sites.google.com/view/debajyotikarmaker/research/preset-path

## References

Schiffner, I., Srinivasan, M.V.. Budgerigar flight in a varying environment: flight at distinct speeds? Biology Letters 2016;12(6):20160221. doi:10.1098/rsbl.2016.0221.

Williams, C.D., Biewener, A.A.. Pigeons trade efficiency for stability in response to level of challenge during confined flight. Proceedings of the National Academy of Sciences of the United States of America 2015;112:3392–3396. doi:10.1073/pnas.1407298112.

Bhagavatula, P.S., Claudianos, C., Ibbotson, M.R., Srinivasan, M.V.. Optic flow cues guide flight in birds. Current biology: CB 2011;21:1794–1799. doi:10.1016/j.cub.2011.09.009.

Lin, H.T., Ros, I.G., Biewener, A.A.. Through the eyes of a bird: modelling visually guided obstacle flight. Journal of the Royal Society, Interface 2014;11:20140239. doi:10.1098/rsif.2014.0239.

Schiffner, I., Perez, T., Srinivasan, M.V.. Strategies for pre-emptive mid-air collision avoidance in budgerigars. PloS one 2016;11:e0162435. doi:10.1371/journal.pone.0162435.

Bhagavatula, P., Claudianos, C., Ibbotson, M., Srinivasan, M.. Edge detection in landing budgerigars (melopsittacus undulatus). PloS one 2009;4:e7301. doi:10.1371/journal.pone.0007301.

Vo, H.D., Schiffner, I., Srinivasan, M.V.. Anticipatory manoeuvres in bird flight. Scientific reports 2016;6:27591. doi:10.1038/srep27591.

Schiffner, I., Vo, H.D., Bhagavatula, P.S., Srinivasan, M.V.. Minding the gap: in-flight body awareness in birds. Frontiers in Zoology 2014;11(1). doi:10.1186/s12983-014-0064-y.

Karmaker, D., Schiffner, I., Wilson, M., Srinivasan, M.V.. The bird gets caught by the WORM: Tracking multiple deformable objects in noisy environments using weight ORdered logic maps. In: Advances in Visual Computing. Springer International Publishing; 2018, p. 332–343. doi:10.1007/978-3-030-03801-4_30.

Wobbrock, J.O., Findlater, L., Gergle, D., Higgins, J.J.. The aligned rank transform for nonparametric factorial analyses using only anova procedures. In: Proceedings of the SIGCHI Conference on Human Factors in Computing Systems. CHI ’11; New York, NY, USA: ACM. ISBN 978-1-4503-0228-9; 2011, p. 143–146. URL: http://doi.acm.org/10.1145/1978942.1978963. doi:10.1145/1978942.1978963.

Batschelet, E.. Circular statistics in biology. @Mathematics in biology; London [u.a.]: Academic Press; 1981. ISBN 0120810506. Literaturverz. S. 353–366.

Biro, D., Meade, J., Guilford, T.. Familiar route loyalty implies visual pilotage in the homing pigeon. Proceedings of the National Academy of Sciences of the United States of America 2004;101:17440–17443. doi:10.1073/pnas.0406984101.

Meade, J., Biro, D., Guilford, T.. Homing pigeons develop local route stereotypy. Proceedings Biological sciences 2005;272:17–23. doi:10.1098/rspb.2004.2873.

Altshuler, D.L., Srinivasan, M.V.. Comparison of visually guided flight in insects and birds. Frontiers in neuroscience 2018;12:157. doi:10.3389/fnins.2018.00157.

